# Fibroblast depletion reveals mammalian epithelial resilience across neonatal and adult stages

**DOI:** 10.1101/2025.07.29.667423

**Authors:** Isabella M. Gaeta, Shuangshuang Du, Clémentine Villeneuve, David G. Gonzalez, Catherine Matte-Martone, Smirthy Ganesan, Deandra Simpson, Hilldana Tibebu, Jessica L Moore, Chen Yuan Kam, Sara Gallini, Haoyang Wei, Fabien Bertillot, Dagmar Zeuschner, Lauren E. Gonzalez, Ushnish Rana, Kaelyn D Sumigray, Sara A Wickström, Valentina Greco

**Affiliations:** Department of Genetics, Yale School of Medicine, New Haven, CT 06510 USA; Department of Cell and Tissue Dynamics, Max Planck Institute for Molecular Biomedicine, 48149 Münster, Germany; Electron Microscopy Facility, Max Planck Institute for Molecular Biomedicine, 48149, Muenster, Germany; Stem Cells and Metabolism Research Program, Faculty of Medicine, University of Helsinki, 00290 Helsinki, Finland; Departments of Cell Biology and Dermatology, Yale Stem Cell Center, Yale Cancer Center, Yale School of Medicine, New Haven, CT 06510 USA; Howard Hughes Medical Institute (HHMI), Chevy Chase, MD, USA

## Abstract

Regenerative organs, like the skin, depend on niche-stem cell interactions that sustain continuous cellular turnover. In cell culture, skin fibroblasts promote epidermal stem cell proliferation and differentiation. Yet, it remains elusive how fibroblasts regulate epidermal stem cell behaviors and differentiation in skin *in vivo*. Here, we asked how fibroblast depletion may impact epidermal stem cell proliferation in the context of adult homeostasis. Surprisingly, we find that significant depletion of fibroblast density does not affect epidermal stem cell proliferative capacity during adult stages *in vivo*. We next probed earlier neonatal stages when skin is actively remodeling but found no change in epidermal stem cell proliferative capacity following fibroblast depletion. These results demonstrate that across different ages, epidermal stem cell proliferative capacity can persist in the face of a largely reduced fibroblast population. Interestingly, neonatal fibroblast depletion does not significantly reduce their secreted collagen I density but affects basement membrane mechanics and epidermal stem cell delamination. Despite these changes, the skin continues to maintain its protective barrier function. Thus, our work demonstrates the skin regenerative program employs robust compensatory mechanisms in the face of fibroblast depletion to maintain functional capacity.

## Introduction

Regenerative organs undergo continuous cell turnover, which involves the production of new cells to replace those that are lost in a niche-regulated manner^1-3^. Skin is an ideal model system to study epithelial regenerative capacity in relationship to its mesenchymal niche because of its continual turnover over the lifetime of an organism and its unique accessibility. Indeed, mechanical changes in the niche environment have been associated with stem cell ageing in the skin and central nervous system^4,5^. In the mammalian skin epidermis, new cells are generated within the epidermal basal stem cell layer by proliferation and move upward by delamination to overlying epithelial layers that maintain the skin barrier. Proper regulation of proliferation and delamination is crucial for maintaining cell density and skin function during homeostasis. The epidermis is in contact with an underlying dermal layer, which is rich in mesenchymal fibroblasts^6^. Co-culture studies demonstrate fibroblasts act as a niche to promote epidermal stem cell proliferation and differentiation^7-10^. Additionally, in skin appendages such as the hair follicle, specialized dermal papilla fibroblasts secrete factors that activate hair follicle epithelial stem cell proliferation^11,12^ and are required for hair follicle growth^13^ and specification of hair epithelial cell type and size^14^. However, how fibroblasts support proliferative demands and differentiation of epithelial cells residing in the epidermal stem cell layer is not understood *in vivo*.

Here, we investigate the relationship between fibroblasts and dynamic turnover in the skin epidermis using mouse models. We asked how fibroblast depletion affects basal cell proliferative capacity during homeostasis in adult mice and demonstrate that fibroblast density depletion *in vivo* does not lead to changes in proliferation in the epidermal stem cell layer. Similarly, fibroblast depletion at neonatal stages - a stage of organ expansion and maturation - also does not affect basal cell proliferative capacity. We find that the major structural component secreted by fibroblasts, collagen I, is not significantly reduced following fibroblast depletion during neonatal stages, however basement membrane (BM) mechanics are perturbed. Moreover, while we find a mild reduction in basal cell delamination, the barrier-protective function of the skin epidermis remains intact following fibroblast depletion during neonatal stages. Together, this study reveals that epidermal basal cell proliferation persists in response to fibroblast depletion, suggesting that the skin employs mechanisms to compensate for fibroblast loss during both skin maturation and homeostasis.

## Results

### Epidermal basal cell proliferation persists despite fibroblast depletion during adult homeostasis

In cell culture, fibroblasts promote epidermal stem cell proliferation and expansion^7-10^. To investigate the potential impact of fibroblasts on the behavior and function of epidermal stem cells in live mice, we employed a skin model - the paw - that lacks remodeling associated with skin appendages and allows maximal surface area of the epidermis to be examined^15,16^. To differentially visualize fibroblasts and epidermal cells, we utilized an inducible Cre ER line that is largely expressed in the fibroblast population (PDGFRα-CreER^17^) in combination with a fluorescent reporter (Rosa-lox-stop-lox-membrane-tdTomato-membrane-GFP, abbreviated LSL-mTmG^18^). Fibroblasts, labeled by membrane GFP, are present throughout the dermal space and interface with epidermal cells labeled by membrane tdTomato and organized in a hexagonal appearance (Figure 1A). To investigate the extent to which fibroblasts impact epidermal basal cell proliferation *in vivo*, we reduced fibroblast density in adult mice (8-11 weeks old) using a mouse model expressing diphtheria toxin fragment A (DTA) under the control of PDGFRα-CreER (LSL-DTA^19^ PDGFRα-CreER, referred to as FibDTA) (Figure 1B). To track fibroblasts, we used both a fluorescent membrane (Cre-dependent reporter either LSL-mTmG or LSL-tdTomato^20^) as well as a nuclear reporter (Cre-independent PDGFRα-H2BGFP^21^). We showed first that PDGFRα-CreER and PDGFRα-H2BGFP label equivalent populations (Supplementary Figure 1A-C). Second, through administration of 100 ug/g tamoxifen for three consecutive days, we detected a ∼70% depletion of fibroblast density in adult fibroblast depleted mice one week post induction with tamoxifen compared to control littermates by imaging of fibroblast nuclei (quantification reflects fibroblast in closest proximity to the epidermis in the top 10 μm of the dremis Figure 1C and E) and corroborated by FACS (Supplementary Figure 1D). Additional marker analyses showed that fibroblast depletion was not specific to a particular subtype (CD26^+^ Sca1^-^, CD26^-^Sca1^+^ or CD26^+^Sca1^+^; Supplementary Figure 1 E,F).

**Figure 1.**
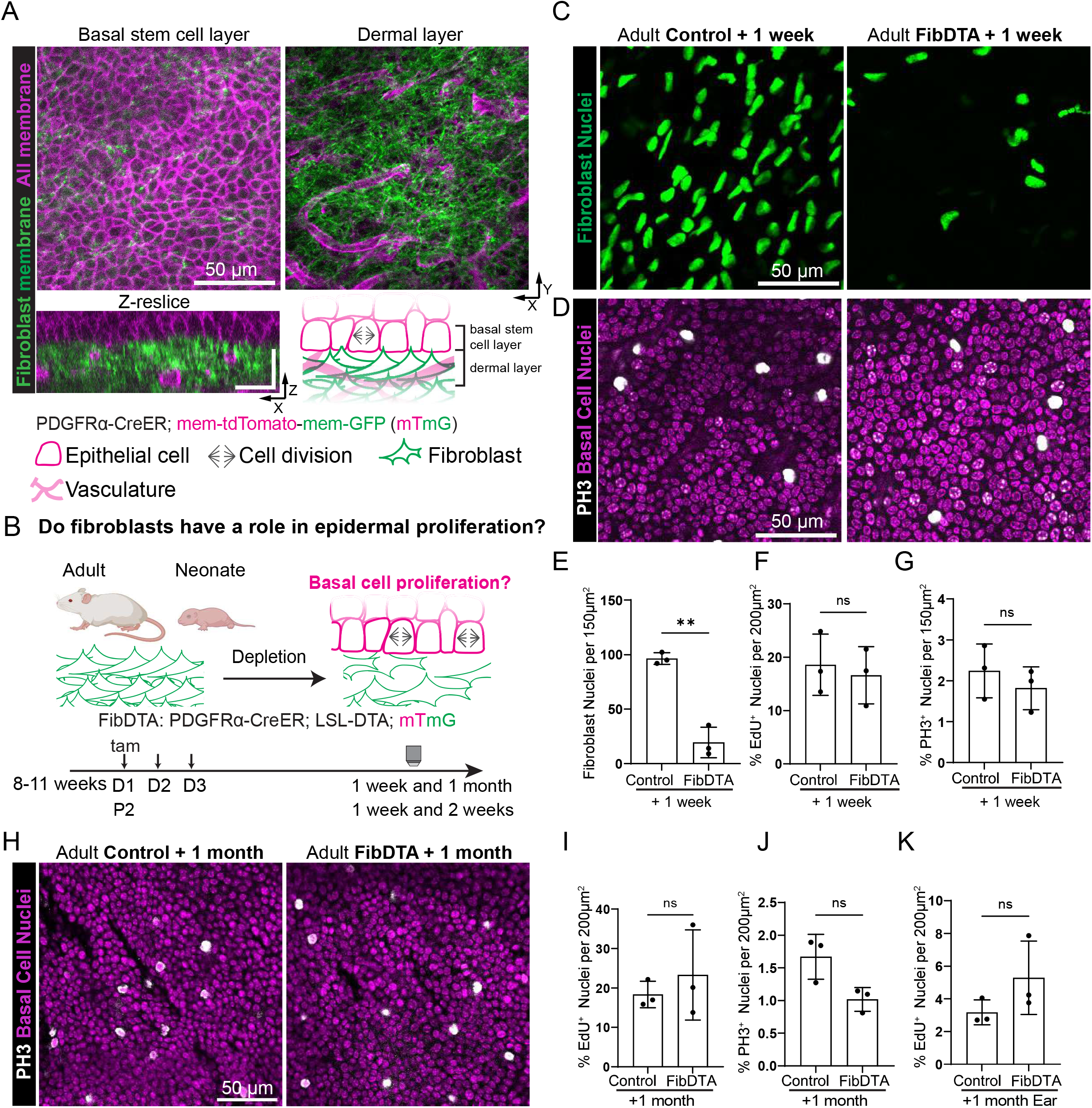
Epidermal basal cell proliferation persists despite fibroblast depletion during adult homeostasis. (A) Live imaging of the non-hairy paw skin of PDGFRα-CreER; LSL-mTmG mice at P10 with fibroblast membrane GFP in green and all cell membrane labeled by tdTomato in magenta. Top left panel: max projection of the epidermal basal stem cell layer 6 μm above the dermal second harmonic generation (SHG) signal as defined in Imaris. Top right panel: max projection of signal 10 μm into the dermal layer as defined in Imaris. Scale bar = 50 μm. Bottom left panel: resliced image of top panels with 1 μm z steps. Scale bars = 25 μm in xy and xz. Bottom right panel: cartoon representation of the proximity between the basal epidermal basal stem cell layer and dermal fibroblasts with their designated icons shown at the bottom. (B) Schematic of the FibDTA model in adult and neonatal mice (PDGFRα-CreER; LSL-DTA; LSL-mTmG). Tamoxifen was given at 8-11 weeks for three consecutive days (D) for adults and P2 for neonates (Figure 2) to induce fibroblast density depletion. Tissue collection or two-photon imaging of the paw or ear skin was done at one week or one month post induction for adults and one week or two weeks post induction for neonates. (C) Representative whole mount images of adult fibroblast nuclei (PDGFRα-H2BGFP) signal in green. Max projection of 10 μm into the dermis of control and FibDTA mice at one week post induction. Scale bar = 50 μm (D) Representative whole mount images of epidermal basal cells stained with PH3 in white and nuclei in magenta from control and FibDTA adult mice one week post induction. Scale bar = 50 μm. (E) Quantification of fibroblast number identified by the PDGFRα-H2BGFP signal in control and FibDTA mice in adult mice one week post induction. Graph represents the average number of fibroblast nuclei in the upper dermal layer (10 μm into the dermis) within a 150 μm^2^ area with SD from n = 3 control and 3 FibDTA mice. ^**^p = 0.0053, unpaired, two-sided, Welch’s t-test. (F) Quantification of %EdU^+^ epidermal basal cells in control and FibDTA adult mice one week post induction within a 200μm^2^ area. n = 3 control and 3 FibDTA mice. p = 0.6875, ns, unpaired, two-sided, Welch’s t-test (G) Quantification of %PH3^+^ epidermal basal cells in control and FibDTA adult mice one week post induction within a 150 μm^2^ area with SD from n = 3 control and 3 FibDTA mice. p = 0.4337, ns, unpaired, two-sided, Welch’s t-test. (H) Representative whole mount images of epidermal basal cells stained for PH3 in white and nuclei in magenta from control and FibDTA adult mice 1 month post induction. Scale bar = 50 μm. (I) Quantification of %EdU^+^ epidermal basal cells in control and FibDTA adult mice 1 month post induction within a 200μm^2^ area. n = 3 control and FibDTA mice. p = 0.1579, ns, unpaired, two-sided, Welch’s t-test. (J) Quantification of %PH3^+^ epidermal basal cells in control and FibDTA mice 1 month post induction. Graph represents the average number of PH3^+^ cells within a 200μm^2^ area. p = 0.0613, ns, unpaired, two-sided, Welch’s t-test. (K) Quantification of %EdU^+^ epidermal basal cells in control and FibDTA adult mouse ear skin 1 month post induction within a 200μm^2^ area. n = 3 control and 3 FibDTA mice. p = 0.2391, ns, unpaired, two-sided, Welch’s t-test.

To investigate the impact of fibroblast depletion on epidermal basal cell proliferation, we performed a pulse of 5-ethynyl-2’-deoxyuridine (EdU) followed by a 6-hour chase to detect cells that have gone through S phase. Remarkably, we found that in the epidermal stem cell layer the number of cells undergoing DNA synthesis per total number of basal cells was similar when comparing fibroblast depleted mice to controls (Figure 1F and Supplementary Figure 1G). Moreover, as EdU incorporation marks broader phases of the cell cycle, we more specifically measured cells undergoing proliferation in M phase by phospho-histone H3 staining (PH3). We found that the percentage of total basal cells undergoing mitotic events does not change in fibroblast depleted mice, nor do the density of the basal cells when compared to control mice (Figure 1D and G, Supplementary Figure 1H). We reasoned that basal cells may demonstrate decreased capacity to proliferate at longer time points following fibroblast depletion. To test this, we induced mice at 11 weeks and collected tissues 1 month post induction. We find that even one month after induction the number of cells undergoing DNA synthesis and mitotic events per total basal cells is not significantly different in fibroblast depleted mice compared to controls (Figure 1H-J, Supplementary Figure 1I). Lastly, EdU assays showed similar results when analyzing the ear, a different geographic skin region, which contains hair follicle appendages (Figure 1K and Supplementary Figure 1J). These findings demonstrate that adult epidermal basal cells retain the capacity to proliferate despite significant fibroblast depletion during adult homeostasis.

### Epidermal basal cell proliferation persists despite fibroblast depletion during neonatal development

We reasoned that during adulthood epidermal stem cells and their niche may have established multiple components to support epithelial proliferation. This prompted us to ask whether proliferative capacity of epidermal stem cells is more reliant on fibroblasts during neonatal development, when the organ is both expanding and maturing to reach adult size. To mitigate systemic effects of fibroblast depletion we began by locally inducing fibroblast depletion in the skin of neonatal mice through topical administration of 4-hydroxy tamoxifen (4-OHT). Administration of 20mg/mL 4-OHT directly to paw skin for two consecutive days (P2 and P3) resulted in a ∼30% depletion of fibroblast density (quantification reflects fibroblast density in the top 10μm of the dermis) and no change in epidermal basal cell density one week post 4-OHT administration (Supplementary Figure 2A-D). To increase the extent of fibroblast depletion similar to experiments in adult mice, we turned to systemic delivery of 167 μg/g tamoxifen at P2, achieving a ∼60% depletion of fibroblast density after 8 days (referred here as 1 week post induction) across subtypes measured by live imaging and corroborated by FACS (Figure 2A and C, Supplementary Figure 2 E-G). Quantification of fibroblast membrane coverage (LSL-DTA; PDGFRα-CreER; LSL-mTmG) demonstrated a ∼10% decrease in fibroblast depleted mice compared to controls (Figure 2B and D). Interestingly, we find that the average nuclei area of remaining fibroblasts throughout the dermis is significantly increased following fibroblast depletion (Figure 2E). We note that systemic depletion of fibroblasts leads to significantly reduced weight of fibroblast depleted neonatal mice (Supplementary Figure 2H).

**Figure 2.**
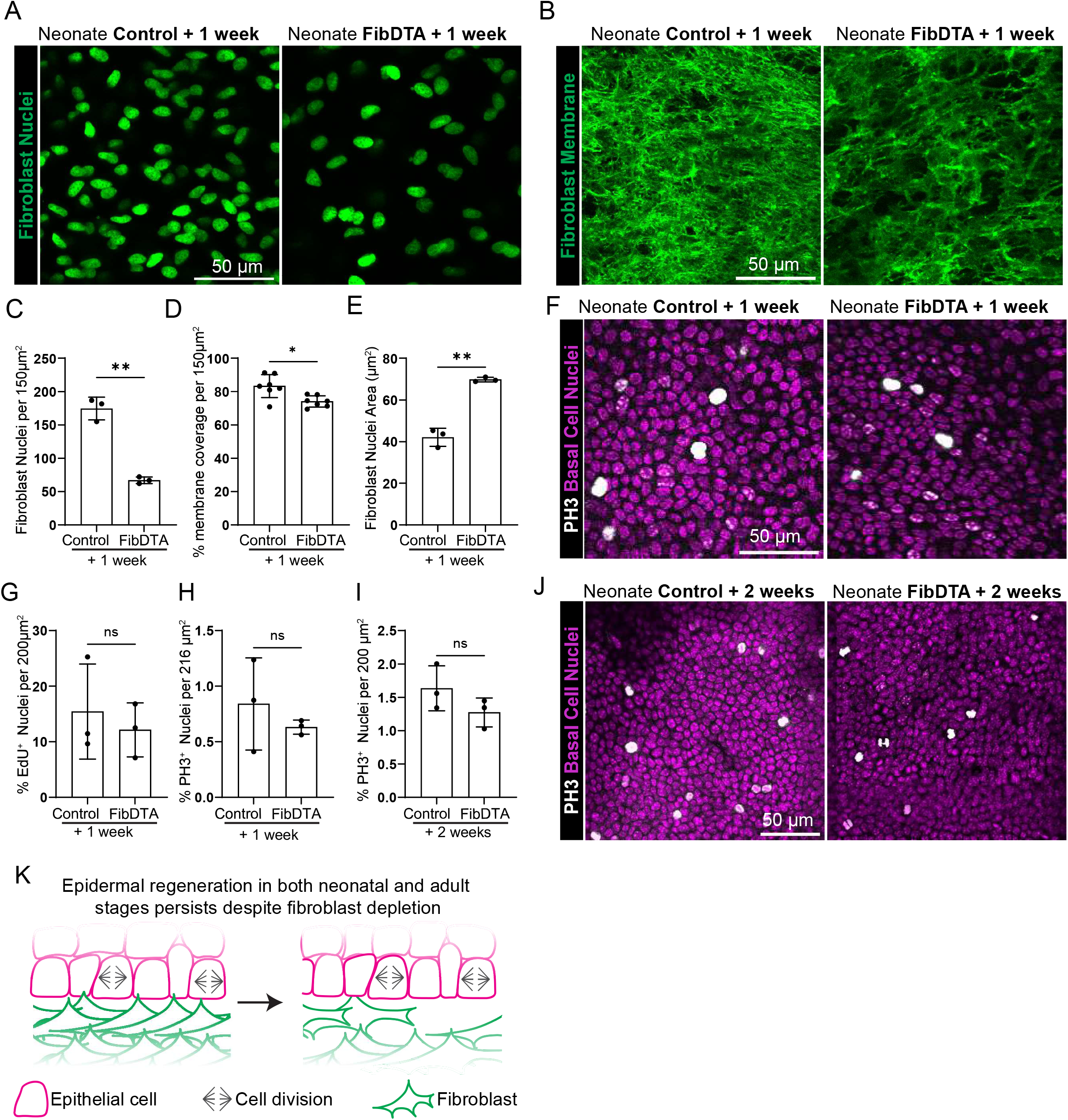
Epidermal basal cell proliferation persists despite fibroblast depletion during neonatal development. (A) Representative neonatal fibroblast nuclei (PDGFRα-H2BGFP) signal in green from live mouse imaging. Max projection of 10 μm into the dermal layer of control and FibDTA neonatal mice one week post induction. Scale Bar = 50 μm. (B) Max projection of fibroblast membrane signal from the upper dermal layer of live mice (10 μm into the dermis) from control and FibDTA neonatal mice one week post induction. Scale Bar = 50 μm. (C) Quantification of fibroblast number identified by the PDGFRα -H2BGFP signal in control and FibDTA neonatal mice one week post induction. Graph represents the average number of fibroblast nuclei in the upper dermal layer (10 μm into the dermis) within a 150 μm^2^ area with SD from n= 3 control and 3 FibDTA mice. ^**^p = 0.0051, unpaired, two-sided, Welch’s t -test. (D) Quantification of the fibroblast membrane coverage in control and FibDTA neonatal mice one week post induction. Graph represents the percent coverage of membrane GFP signal max projection in the upper dermal layer (10 μm into the dermis) within a 150 μm^2^ area with SD from n = 7 control mice and 7 FibDTA mice. *p = 0.0115, unpaired, two-sided, Welch’s t t-test. (E) Quantification of fibroblast nuclei area identified by the PDGFRα -H2BGFP signal in control and FibDTA neonatal mice one week post induction. Graph represents the mean of fibroblast nuclei throughout the dermis within a 150 μm ^2^ area with SD from 414 nuclei in n = 3 control and 155 nuclei from n= 3 FibDTA mice. (F) Representative whole mount images of epidermal basal cells stained with PH3 in white and nuclei in magenta from control and FibDTA neonatal mice one week post induction. The isolation of the epidermal basal cell layer relative to the other layers was determined by the small and compact nuclei of epidermal basal cells stained with hoechst and their proximity to SHG signal. Scale bar = 50 μm. (G) Quantification of %EdU^+^ epidermal basal cells in control and FibDTA neonatal mice one week post induction within a 200μm^2^ area. n = 3 control and 3 FibDTA mice. p= 0.5984, ns, unpaired, two-sided, Welch’s t-test. (H) Quantification of %PH3^+^ epidermal basal cells in control and FibDTA mice one week post induction. Graph represents the average number of PH3^+^ cells within a 216 μm^2^ area with SD from n = 3 control and 3 FibDTA mice. p = 0.4745, ns, unpaired, two-sided, Welch’s t-test. (I) Quantification of %PH3^+^ epidermal basal cells in control and FibDTA mice two weeks post induction within a 200 μm^2^ area with SD from n = 3 control and n = 3 FibDTA mice. p = 0.2047, ns, unpaired, two-sided, Welch’s t-test. (J) Representative whole mount images of epidermal basal cells stained with PH3 in white and nuclei in magenta from control and FibDTA mice two weeks post induction. Scale bar = 50 μm. (K) Schematic summary of the impact of fibroblast depletion on epidermal stem cell regeneration in neonatal and adult mice.

To determine whether systemic fibroblast depletion impacts epidermal basal cell proliferation at neonatal stages, we again measured DNA synthesis and mitotic events by using a 6-hour EdU chase and PH3 staining respectively. We found that the number of cells undergoing DNA synthesis per total basal cells was similar when comparing fibroblast depleted and control neonatal mice one week post induction (Figure 2G and Supplementary Figure 2I). Additionally, we find that the percentage of total basal cells undergoing mitotic events also does not change between fibroblast depleted and control neonatal mice (Figure 2F and H). We find the same trend in epidermal basal cells undergoing mitotic events when comparing fibroblast depleted and control neonatal mice two weeks post induction (Figure 2 I,J).

Large population depletion can cause immune reactions^22^, which may also be contributing as a compensatory mechanism to the sustainment of epidermal proliferation. We showed that our DTA model does not change macrophage, neutrophil and monocyte abundance, and led to a 1% increase in T-cells locally to paw tissues (Supplementary Figure 3 A-D). Furthermore, we did not detect an increase in circulating cytokine levels in serum (Supplementary Figure 3 E). Additionally, we did not detect gross architectural changes to vasculature diameter in the nearby capillary plexus upon fibroblast depletion (Supplementary Figure 3 F and G).

Taken together, these findings demonstrate that epidermal basal cells retain their capacity for proliferation upon fibroblast depletion in neonatal contexts, paralleling the results obtained in the adult context (Figure 2K). These results suggest that multiple sources support epidermal stem cell growth upon fibroblast loss *in vivo*.

### Neonatal fibroblast depletion alters basement membrane mechanics and epidermal delamination with no impact on collagen density or tissue function

Fibroblasts have a major role in producing and remodeling the extracellular matrix, including the basement membrane^6,23^. We assessed how collagen I, the major structural component of the dermal extracellular matrix, and the basement membrane, the substrate to which the epithelium is anchored, may be altered. Surprisingly, fibroblast depletion in neonatal mice 1 week post induction does not lead to a statistically significant reduction in collagen I density as measured by second harmonic generation (SHG) mean fluorescence intensity (MFI; quantification reflects MFI in the top 10 μm of the dermis; Figure 3A and B). Furthermore, we measured collagen fiber thickness and lacunarity (measurement of how gaps in signal are distributed through an image) to determine whether the spatial patterning of collagen fibers differs between fibroblast depleted and control mice, and find no significant difference (Figure 3C and D). These results suggest robust compensatory mechanisms maintain dermal structural integrity. Next, we interrogated basement membrane morphology and mechanics in fibroblast depleted neonatal mice. Morphologically, we find that the basement membrane is still intact by transmission electron microscopy when comparing fibroblast depleted mice to controls (Figure 3E). To interrogate whether basement membrane mechanics may be perturbed despite appearing morphologically intact, we turned to atomic force microscopy (AFM) to measure basement membrane stiffness in frozen tissue sections (Figure 3F) and found significantly reduced basement membrane stiffness in fibroblast depleted mice compared to controls (Figure 3G).

**Figure 3.**
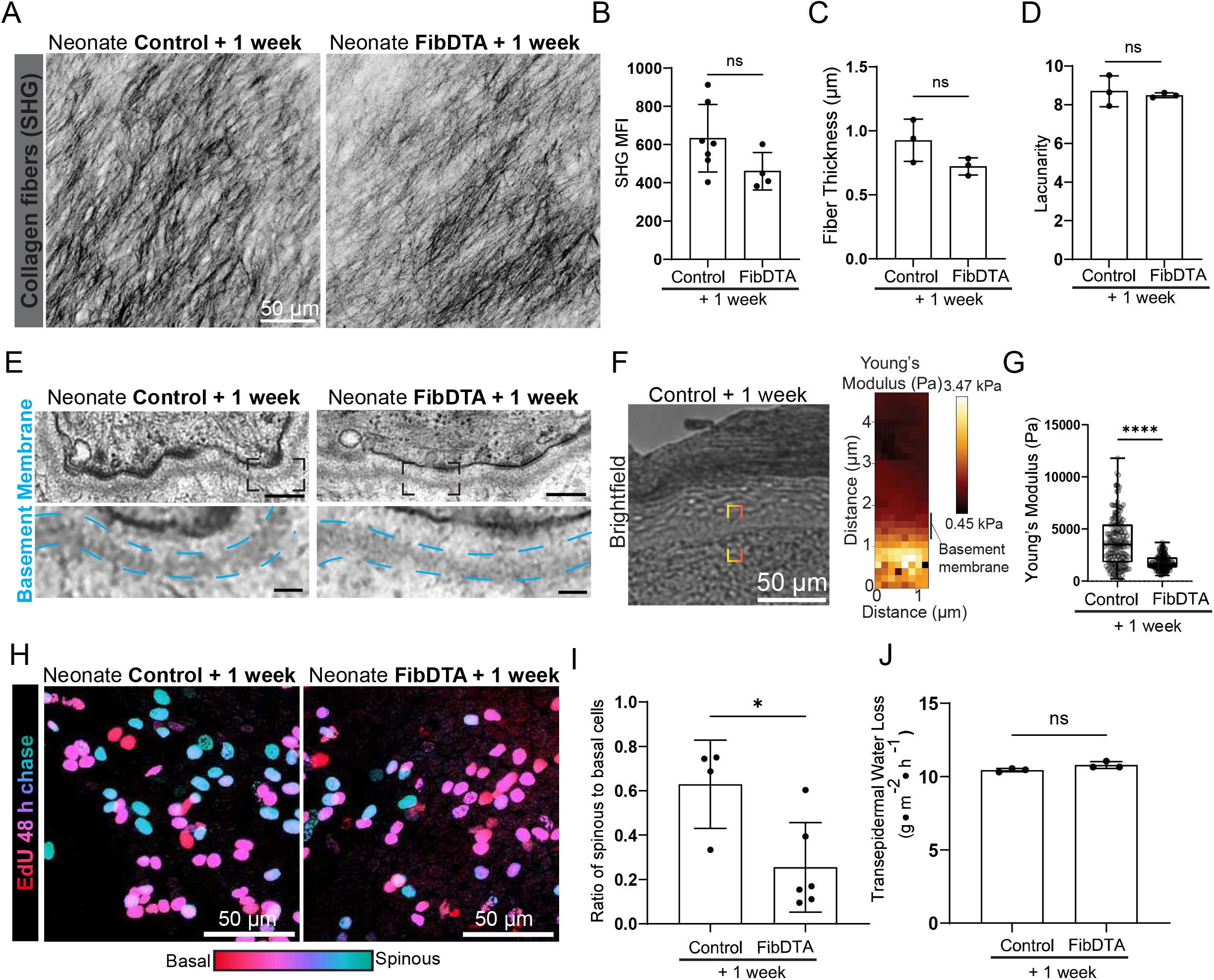
Neonatal fibroblast depletion alters basement membrane mechanics and epidermal delamination but not collagen density or tissue function. (A) Representative max projection of the SHG signal within 10 μm into the dermis of live mice, revealing the structure of collagen fibers from control and FibDTA neonatal mice one week post induction. Scale bar = 50 μm. (B) Quantification of the collagen fiber density in the upper 10 μm of the dermis in control and FibDTA neonatal mice one week post induction. Graph represents average MFI of SHG signal with SD from 7 control mice and n = 4 FibDTA mice. p = 0.0674, ns, unpaired, two-sided, Welch’s t-test. To address unequal group sizes, we performed 100 random downsampling iterations (control subsampled to n=4). These results confirmed a consistent trend (mean difference = 179.214, 95% CI = 60.228,279.408), but did not reach statistical significance (mean p value = 0.2034, iterations with p<0.05 = 5/100). (C) Quantification of fiber thickness in control and FibDTA neonatal mice one week post induction. Graph represents mean thickness with SD from from n = 3 control and 3 FibDTA mice. p = 0.1338, ns, unpaired, two-sided, Welch’s t-test. (D) Quantification of fiber lacunarity in control and FibDTA neonatal mice one week post induction. Graph represents mean lacunarity with SD from n = 3 control and 3 FibDTA mice. p = 0.6888, ns, unpaired, two-sided, Welch’s t-test. (E) Representative transmission electron micrographs of basement membrane in control and FibDTA neonatal mice one week post induction. Dashed boxes indicate zoom regions to highlight basement membrane, outlined in cyan. Top box scale bar = 200 nm, zoom box scale bar = 50nm. (F) Representative brightfield image produced during AFM measurements and corresponding heat map of young’s modulus to measure basement membrane stiffness. Red dashed box indicates representative region of young’s modulus heat map measurement. Scale bar = 50 μm (G) Box and whiskers plot of Young’s elastic moduli from atomic force indentation spectroscopy measurements in skin BM regions from control and FibDTA mice neonatal mice one week post induction. Outliers were identified using the ROUT method and removed, resulting in n= 147 force curves from 3 control (LSL-DTA or Col1a1 fl/fl) mice and n = 122 force curves from 3 FibDTA mice. ****p = <0.0001, Kolmogorov-Smirnov test. (H) Images showing EdU whole mount staining after 48-hour pulse-chase coded by depth with red (epidermal basal cells) closer to the dermis and blue (epidermal spinous cells) further into the epidermis from control and FibDTA neonatal mice one week post induction. Relative locations of EdU^+^ cells to the epidermal basal cell layer are noted, and cells are categorized as EdU^+^ spinous cells versus EdU^+^ basal cells. Scale bars = 50 μm. (I) Quantification of the rate of delamination in control and FibDTA neonatal mice one week post induction. Graph represents the average ratio of EdU^+^ spinous to basal cells with SD from n = 4 control and 6 FibDTA mice. ^*^p = 0.0244, unpaired, two-sided, Welch’s t-test. (J) Quantification of transepidermal water loss measurements between control and FibDTA mice at neonatal mice one week post induction. n = 3 control and 3 FibDTA mice. p = 0.1007, ns, unpaired, two-sided, Welch’s t-test.

As the epidermal stem cell layer is anchored to the basement membrane, we next asked whether fibroblast depletion could impact basal cell delamination. To examine this, we performed a 48-hour EdU chase experiment to track EdU^+^ differentiating basal cells that have reached the overlaying differentiated layer, the spinous layer, in fibroblast depleted mice compared to controls. A ratio of EdU^+^ cells in the spinous versus basal cell layer was used to estimate delamination from the basal layer. We find decreased delamination of differentiating epidermal basal cells in fibroblast depleted mice compared to controls (Figure 3H and I). We also performed immunofluorescence on neonate paw samples 1 week post induction for stemness marker K14 and differentiation marker K10 and quantified the percentage of K10^+^ in the epidermal basal layer (Supplementary Figure 2 J and K). We found no significant difference in K10^+^ basal cells between fibroblast depleted and control mice. These changes in basement membrane stiffness and differentiating epidermal basal cell delamination prompted us to assess overall barrier function by measuring transepidermal water loss (TEWL). We found no significant difference in TEWL measurements between fibroblast depleted mice compared to controls, indicating proper epidermal barrier function is maintained (Figure 3J).

Taken together, despite perturbations to basement membrane stiffness and epidermal basal cell delamination, neonatal dermal collagen density and epithelial barrier function are retained in the face of significant fibroblast depletion.

Fibroblasts are major contributors of organ integrity including through both secretion of critical structural elements and their ability to influence regeneration of neighboring epidermal stem cells^6,24,25^. Our results demonstrate that epidermal basal cell proliferation persists despite significant fibroblast depletion during both neonatal and adult stages, suggesting that the program to support epidermal regeneration is built in excess. These results parallel hair follicle studies which demonstrate that a critical number of dermal papilla fibroblasts are necessary to promote hair follicle epithelial stem cell growth^14^. This is not the case under conditions where the entire population of dermal papilla fibroblasts have been eliminated, which inhibits hair follicle growth entirely^13^. Moreover, reorganization of dermal papilla architecture leads to shorter hair production but still supports hair follicle stem cell proliferation and differentiation^26^. These results demonstrate that maintained presence of fibroblasts can still support stem cell growth even if architecture is altered.

In our system, a fraction of dermal fibroblasts remains after depletion, and epidermal basal cell proliferation continues, unmasking the potential for fibroblast compensation and redundancy in the dermis. This compensation is evident as neonatal fibroblast depletion leads to an increase in fibroblast nuclei area, suggesting fibroblasts may modulate their cell size upon loss of their neighbors. Indeed, the change in fibroblast nuclei area in response to depletion may mimic changes in intestinal epithelium nuclei size in response to feeding and starvation^27^. Interestingly, while there is a significant decrease in fibroblast membrane coverage, this decrease does not reflect the depletion from counting fibroblast nuclei (10% membrane coverage reduction vs. 60% nuclei reduction) suggesting either that remaining fibroblasts hold the capacity to expand their membrane area or that the tissue is already built with intertwined architecture that can largely remain after depletion. Indeed, in aged mice, fibroblasts are lost in clusters, and also demonstrate membrane coverage compensation^15^. Maintenance of fibroblast membrane coverage may also aid the observed maintained collagen density between neonatal control and fibroblast depleted conditions. It is conceivable maintained membrane coverage ensures secretion is equally spread and that remaining fibroblasts augment their collagen I synthetic capacity to provide similar output generated by a larger fibroblast population.

Furthermore, results in this study reflect remarkable tissue capacity for compensation for critical physiological processes of other organ systems. For instance, intestinal brush border microvilli, which are essential for nutrient absorption, are supported by actin filaments and stabilized by actin bundling proteins. Remarkably, microvilli can still form despite knockout of three of the four known actin bundling proteins^28,29^, revealing a striking example of molecular redundancy to maintain tissue integrity.

Our study also complements observations of epidermal stem cell growth and differentiation in cell culture. These cell culture studies demonstrate a greater dependence of epidermal stem cells on fibroblasts for differentiation, whereas proliferation of epidermal stem cells is enhanced by the presence of fibroblasts but can continue over short periods or at high epidermal seeding densities in the absence of fibroblasts^7-10^. While our observations suggest that delamination may be slowed down, the proportion of K10^+^ basal cells committed to delaminate is not significantly changed.

Our study examines epidermal proliferative capacity in short and longer time points following fibroblast depletion in both neonatal and adult mice, demonstrating that the epidermal stem cell layer retains the capacity to proliferate at both stages. Future studies should aim to understand whether the full repertoire of fibroblasts is necessary to sustain epidermal basal cell proliferation under wounding conditions following fibroblast depletion.

Overall, this work has important implications for understanding the regulation of epidermal stem cell behaviors in relation to dermal fibroblasts in live mice. Our work suggests that niche dependence may be consistent in skin appendages versus epidermis, underscoring the need for further comprehensive understanding of the complex interplay between different cell types in a physiological microenvironment.

## Methods

### Mice

The following mice were procured from the Jackson Laboratory: LSL-DTA (#009669)^19^, PDGFRα-CreER (#018280)^17^, LSL-mTmG (#037456)^18^, PDGFRα-H2BGFP (#007669)^21^ and LSL-tdTomato^20^ (#007909). FibDTA mice carried both PDGFRα-CreER and LSL-DTA alleles. Depending on requirement for cell labeling and mouse availability, FibDTA mice additionally carried one or more fluorescent reporters (LSL-mTmG, PDGFR -H2B-GFP, LSL-tdTomato). Control mice carried either PDGFRα-CreER or LSL-DTA and one or more fluorescent reporters depending on the requirement for cell labeling and mouse availability, except where indicated in the figure legend, which included Col1a1^flox/flox^ mice without PDGFRa-CreER, wherein loxp sites flank exons 2-5 of the COL1A1 (Mayumi Ito, New York University). Adult mice were defined as mice 8-11 weeks of age at the time of induction, with imaging or tissue harvesting occurring 7 days or 30 days after the first day of induction. Neonatal mice were defined as post-natal day 2 (P2) at the time of induction, with imaging or tissue harvesting occurring 8 days (referred to as 1 week) or 14 days after induction. At the indicated time points, mice from experimental and control groups were littermates selected at random – independent of sex – for tissue collection. No blinding was done. All procedures involving animal subjects were performed under the approval of the Institutional Animal Care and Use Committee (IACUC) of the Yale School of Medicine.

### Induction with Tamoxifen and 4-Hydroxytamoxifen

To induce systemic Cre recombination in neonatal mice, 50 μL of 6.6 mg/mL tamoxifen (167 μg/g) was administered intragastrically at P2. Mice from experimental and control groups were selected at random – independent of sex – at P10 and used for live imaging and tissue collection. To induce Cre recombination in adult mice 100 μg/g tamoxifen was administered for three consecutive days to 8 11-week-old mice intraperitoneally. Topical administration of 4-hydroxy tamoxifen (4-OHT) was conducted by first anaesthetizing mice using 2% isoflurane in oxygen and air and directly pipetting 20mg/mL of 4-OHT dissolved in DMSO to paw skin for 30 minutes. This procedure was repeated in neonatal mice for two consecutive days (P2 and P3).

### *In vivo* imaging

Live imaging of mice was performed on the non-hairy, flat region of the right hind paws. Mice were anaesthetized by inhaling 2% isoflurane in oxygen and air. The hind paw was mounted on a custom-made stage with a warming pad and the mice remained anaesthetized with 1.5% isoflurane during imaging. A glass coverslip was placed against the region of interest with a water interface to minimize the vibration of breathing. Images were acquired with a LaVision Biotec (Miltenyi Biotec) two-photon microscope equipped with two tunable Ti:Sapphire lasers, a Chamelon Vision II (Coherent) (940 nm) and a Chameleon Discovery (Coherent) (1120 nm) through a 40X water immersion lens (NA 1.1; Zeiss or NA 1.15; Nikon) in 1 μm Z-steps. Mature collagen fibers capable of second harmonic generation (SHG) were excited by the 940 nm laser to emit signal collected in the blue channel.

### Immunofluorescence

For whole-mount staining, paw skin tissue was isolated and fixed in a 4% paraformaldehyde (PFA) solution in PBS overnight at 4 C or for 4hrs at room temperature. The tissue was then transferred to an Eppendorf tube for blocking with 0.2% Triton X-100, 5% normal donkey serum, and 1% BSA in PBS. The tissue was incubated with primary antibodies for 48 hours at 37 C or over 60 hours at room temperature, washed, and then incubated with secondary antibodies for 24 - 60 hours at room temperature. The primary antibodies used in this study were rabbit anti-pH3 (1:300; Millipore, 06-750), guinea pig anti-keratin 10 (1:200; ARP, 03-GP-K10) and rabbit anti-keratin 14 (1:400; BioLegend 905301). The secondary antibody used in this study were anti rabbit AlexaFluor 647 (1:200; Thermofisher). For nuclei staining, Hoechst 33342 (1:2000; Becton Cickinson: H3570) was added during the secondary incubation or SiR-DNA (1:500; Cytoskeleton inc: SKU CY-SC007) was added overnight after secondary antibody incubation. Lastly, the tissues were mounted with Vectashield Anti-fade mounting medium (Vector Laboratories) or SlowFade Diamond Antifade Mountant (ThermoFisher) with an overlying coverslip sealed with nail polish. The slides were imaged on the LaVision two-photon microscope as described in “*In vivo* imaging” with 1-2 μm Z-steps.

### Electron Microscopy

Paws were cut and directly transferred in 4% formaldehyde, 0.1 M cacodylate buffer, pH 7.4 at 37 C and further dissected to small stripes to allow better infiltration of the fixatives. Fixative was exchanged to 2% formaldehyde, 2% glutaraldehyde, 0.1 M cacodylate buffer, pH 7.4 for 30 min at 37 C and left for another 2 hours at room temperature. The fixed tissue was embedded in LMP agarose and sliced into 250 μm thick sections with a vibratome (VT1200S, Leica, Vienna). These got post fixed in 1 % osmium tetroxide, 1.5 % potassium cyanoferrate for 1 hour, stepwise dehydrated with ethanol, including 0.5 % uranyl acetate en-bloc stain in the 70% ethanol incubation step. Final dehydration was achieved by 2 changes in propylenoxide. The samples were embedded in epon and 60 nm ultrathin sections were cut from the region of interest on an ultramicrotome (UC 6, Leica, Vienna), stained with lead and analysed at a transmission electron microscope (Tecnai 12-biotwin, Thermofisher scientific, the Netherlands). Representative images were acquired with a 2K CCD camera (Veleta, EMSIS, Muenster).

### Atomic Force Microscopy

For Atomic Force Microscopy (AFM), paw skin was dissected, nitrogen frozen and then embedded in OCT blocks. AFM measurements of the basement membrane were performed on freshly cut 20 μm cryosections using JPK Nano Wizard 2 (Bruker Nano) mounted on an Olympus IX73 inverted fluorescent microscopy (Olympus) and operated via JPK SPM Control Software v.5. Cryosections were equilibrated in PBS supplemented with protease inhibitors and measurements were performed within 20 minutes of thawing the samples. Triangular non-conductive Silicon Nitride cantilevers (MLCT, Bruker Daltonics) with a nominal spring constant of 0,07 Nm^-1^ were used for the nanoindentation experiments of the apical surface of cells and nuclei. For all indentation experiments, forces of up to 3 nN were applied, and the velocities of cantilever approach and retraction were kept constant at 10 μm^s-1^ ensuring an indentation depth of 500 nm. All analysis were performed with JPK Data Processing Software (Bruker Nano). Prior to fitting the Hertz model corrected by the tip geometry to obtain Young’s Modulus (Poisson’s ratio of 0.5), the offset was removed from the baseline, contact point was identified, and the cantilever bending was subtracted from all force curves.

### EdU Treatment and Quantification

EdU, 5-ethynyl-2’-deoxyuridine, is a thymidine analog that incorporates into DNA. An EdU pulse chase experiment was adopted to examine the differentiation process of the epidermis and epidermal proliferative capacity^30^. To label basal cells, mice were pulsed with Edu (50 μg/g in PBS) via intraperitoneal injection. For the 48 hour EdU chase, mice were pulsed at P8, and 48 hours after the injection at P10, paw skin tissues were isolated and fixed in 4% aqueous PFA solution overnight. For the 6 hour EdU chase paw and ear skin tissues were isolated and fixed 6 hours after injection at the indicated timepoint in 4% aqueous PFA solution 4 hours at room temperature or overnight at 4 C. EdU incorporation was evaluated by the Click-iT EdU Cell Proliferation Kit for Imaging Alexa Fluor 488 dye (48 hour EdU Chase, Invitrogen C10337) or Alexa Fluor 647 and Alexa Fluor 555 dye (6 hour EdU chase, C10340, C10338) based on the manufacturer’s instruction. The tissues were then stained as described in “whole-mount staining”. After image acquisition, the delamination status of EdU positive cells were expressed as the distance from basal cells manually analyzed in FIJI using the line measurement for the 48 hour EdU chase. For the 6 hour EdU chase, EdU^+^ cells were counted and plotted as a percentage of total basal epidermal stem nuclei.

### Cytokine assay

Lipopolysaccharide (LPS) was injected intraperitoneally at 2 μg/g in CD1 mice at P9 as a positive control for inflammation. Blood samples from P9 mice of respective treatments including controls were collected from the neck at the time of decapitation. The samples were spun-down, and serum was collected and kept at -80°C until the assay was performed. To assess inflammatory cytokines and growth factor cytokines in our models, we performed a Bio-Plex ProTM Mouse Cytokine 8-Plex Assay and Bio-Plex ProTM Mouse Cytokine 9-Plex Assay respectively (Bio-rad laboratories; M60000007A, MD000000EL). The assays were performed according to the manufacturer’s instructions. Briefly, serum samples were thawed and diluted 1:4 with provided diluent. 50 L of each sample including standards and blanks were added to the 10X coupled bead pre-coated plate and incubated for 30 mins on a shaker at room temperature. The samples were washed three times and incubated with detection antibody for an additional 30 minutes on a shaker at room temperature, washed three times and incubated with SA-PE for 10 minutes. The samples were then washed for a final three times and resuspended in 125 L of assay buffer before running on a Luminex 200 analyzer with a dual laser detection platform.

### Fluorescence Activated Cell Sorting

Neonate and adult paws were collected and processed into single cell suspensions using an adapted protocol from Yang et al. (2017, Cell 169, 483 496). Specifically, paws were placed dermis side down in 0.25% collagenase IV (Sigma; C5138) in HBSS (GIBCO;14170-112) for 45 minutes followed by 20 minutes in 0.25% trypsin (GIBCO; X). After digestion, the paws were scraped using a blunt blade to create single cell suspensions and washed in FACS Buffer (3% FBS, 2mM EDTA). The suspensions were then centrifuged at 350G for 10 minutes and filtered through a 40μm filter (Falcon; 352340) before staining. The following antibodies were used for staining. Fibroblast heterogeneity: CD26-PE (Biolegend;137803), Sca-1 APC (Invitrogen;17-5981-83). Immune profiling: CD45-FITC (Biolegend;147710), Ly-6G-PE-efluor™610 (Invitrogen; 61-9668-82), CD11b-APC (Invitrogen; 17-0112-82), CD45-PE (Biolegend;137803), F4-80-APC (Biolegend; 123116), CD3 APC (Biolegend; 100236). Cells were acquired on a Becton Dickinson LSRII Flow Cytometer outfitted with Diva software v 8.0.1, and the data was analyzed using Flowjo v.10.9.0.

### Imaging Analysis

#### Fibroblast quantification and second harmonic generation (SHG) analysis

Raw live imaging data was imported into FIJI (ImageJ, NIH) for stitching^31^. To reduce the dimensionality of 3D imaging stacks we used Imaris software (Oxford Instruments) to isolate the upper dermal signal and a custom MATLAB script (MathWorks) to flatten it. Upon importing the image stack into Imaris, the dermal space was defined by creating a surface from the SHG collagen signal. Using Imaris’ distance transformation tool, a new channel was created to represent distance from the dermal surface. A third surface, an inverse of the original dermal surface, was used to generate a second distance transformation channel representing distance into the dermis. This allowed us to mask a defined distance into the dermis from the epidermal-dermal interface. The resulting masked channels were exported back to FIJI and run through our custom MATLAB script for flattening and generation of maximum intensity projections for further MFI analysis. Consistent thresholding in Fiji was applied to all maximum intensity projections to quantify the relative % membrane coverage based on area. The adult fibroblast nuclei count was conducted using Imaris software. The PDGFRaH2BGFP signal was imported into Imaris as a z-stack. Next, the Imaris spot function was used to identify nuclei spots, with a spot diameter of 7 μm. Any inaccuracies in thresholding were manually corrected in Imaris before exporting statistical data for the final nuclei count. Quantifications of fibroblast nuclei density, fibroblast membrane coverage, SHG MFI, fiber thickness and lacunarity were all confined to the top 10 μm of the dermis. Collagen fiber thickness and lacunarity were calculated using the AnaMorf FIJI plugin. Fiber thickness was defined as (% High density matrix x Total Image Area)/Total Fiber Length. Quantification of nuclei area was conducted on 2D max intensity projections of the whole dermis, with nuclei masks first created using CellPose to generate instance segmentation masks, wherein each detected nucleus was assigned a unique integer label. Morphometric properties of segmented nuclei were extracted using the regionprops and regionprops_table functions from the scikit-image library (skimage.measure). CellPose output masks were cast to 64-bit integer arrays and passed directly to these functions. For each labeled region, cross-sectional area (μm^2^) was computed. Capillary diameter was measured using the VasoMetrics FIJI plugin on 8-16μm flattened projections of the upper dermis.

#### Epidermal nuclei counting

Whole mount image stacks were processed using FIJI to identify distinct layers of cells based on their proximity to the basal layer and the SHG signal in the dermis. Epidermal basal cells were identified based on their small and compact nuclei, and were either counted manually using FIJI’s cell counter or in Imaris by first using the same surface/masking approach described above, except a mask was applied just above the epidermal-dermal interface to isolate the basal layer. Next, the Imaris spot function was used to identify nuclei spots. Any inaccuracies in thresholding were manually corrected in Imaris before exporting statistical data for the final nuclei count. The number of PH3^+^ and EdU^+^ cells were quantified using FIJI’s cell counter, defined by large morphological “bright spots” overlapping with nuclear signal. %PH3^+^ and EdU^+^ cells were calculated by dividing the average number of PH3^+^ or EdU^+^ basal cells per mouse by the average number of total basal cells per mouse.

The %K10^+^ basal cell calculation was determined by counting the total number of cells in the basal layer using FIJI’s cell counter and dividing by the total number of basal cells expressing K10.

### Trans-epidermal water loss measurement

Barrier function was determined using a Tewameter TM Nano (Courage and Khazaka electronic). P10 mice were anaesthetized by inhaling 2% isoflurane in oxygen and air and then probed on the paw to measure trans-epidermal water loss. The data was acquired using the CK-MPA software provided by the manufacturer. The data was then exported to excel for analysis.

### Statistics and reproducibility

Biostatistical analyses were performed with the GraphPad Prism (Version 9.4) software (GraphPad Inc., La Jolla, CA). The distribution of replicates was described by means with standard deviations in the graphs unless indicated. Statistical comparisons between conditions were made using the unpaired two-tailed Welch’s t-test, Kolmogorov-Smirnov test, or one-way ANOVA where indicated. To account for unequal group sizes between control (n=7) and FibDTA (n=4) animals in SHG analysis (Figure 3B), a random downsampling analysis was performed in Python. For each of 100 iterations, four mice were randomly selected without replacement from the control group and compared against all four FibDTA mice using a two-sided Mann-Whitney U test. The mean difference in MFI and corresponding 95% confidence interval were calculated across all iterations. All analyses were performed using the NumPy and SciPy libraries. The random seed was set to 42 to ensure reproducibility.

## Data availability

All the data that support the findings of this study are available from the corresponding author upon reasonable request.

## Acknowledgments

We thank the Greco lab members (specifically Tianchi Xin), thesis committee members of SD Stefania Nicoli, and Erica Herzog), as well as Mayumi Ito for helpful discussions and constructive feedback on the manuscript. We thank former Greco lab members Edward Marsh and Katie Cockburn for mentoring SD on scientific thinking. We thank Ryan Driskell for helpful discussion and expertise in fibroblast biomarkers, as well as Salvador Aznar-Benitah and Guiomar Solanas for helpful discussion. VG is supported by an HHMI Investigator Award, the Leo Foundation (LF-OC-23-001463), the European Research Council (101167365, BEMOSAIC) and the National Institutes of Health (R01AR063663, R01AR072668). HT is supported by the Yale Science, Technology and Research Scholars Program (STARS). The content is solely the responsibility of the authors and does not necessarily represent the official views of the National Institutes of Health. Research in the SW lab is funded by the Max Planck Society, European Research Council (ERC) under the European Union’s Horizon 2020 research and innovation program (grant agreement 770877 - STEMpop) and Academy of Finland Center of Excellence BarrierForce (346132). CV is the recipient of the Marie Curie fellowship H2020-MSCA-IF-101032331.

## Author Contributions

The work presented in this research publication was made possible by the contributions of a dedicated team of researchers. IG, SD and VG collaborated to develop the project and its main conceptual ideas, with VG and SW overseeing the project. Mouse work and the generation of mouse lines were conducted by IG, SD, CM, and HW. IG, SD, S Ganesan, CM, CV, DS, HT and S Gallini planned and carried out the experiments. The interpretation of the results was contributed by IG, SD, S Ganesan, JM, CK, CV, FB, S Gallini, KS, SW, and VG. The analysis was performed by, IG, SD, S Ganesan, DG, JM, DS, HT, UR and FB. IG, SD, SW, and VG wrote the manuscript, with the assistance of LG, CM, JM, CK, CV, and input from all authors. All authors discussed the results and made significant contributions to the final manuscript.

