## Supplementary Figures 1-3 for "Fibroblast depletion reveals mammalian epithelial resilience across neonatal and adult stages"

Supplementary Figure 1

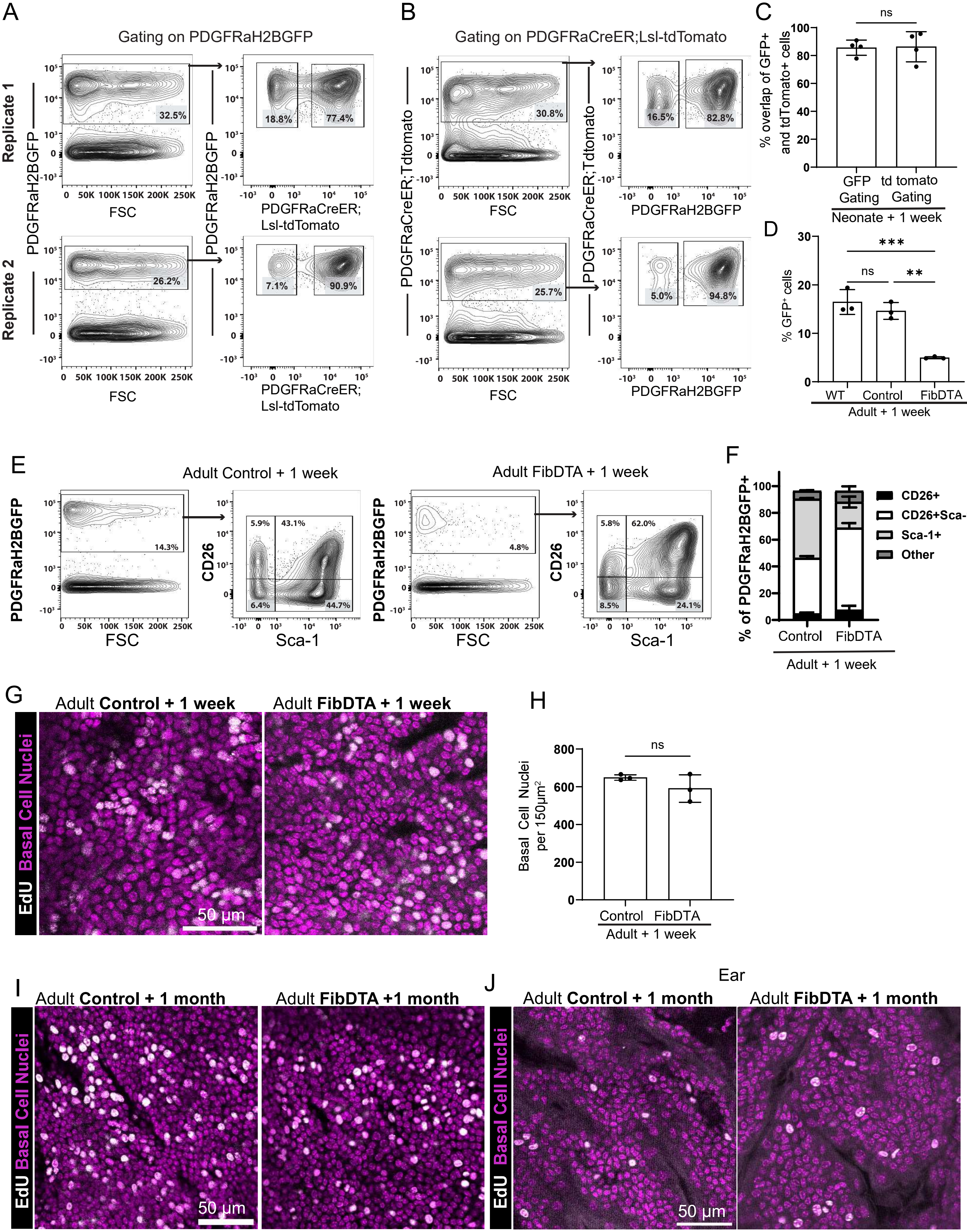

**Supplementary Figure 1.**

(A) Representative FACS plots from two replicate neonatal mice one week post induction. Mice expressing PDGFRα-H2B-GFP; PDGFRα-CreER; lsl-tdTomato gating first on GFP+ cells and then tdTomato+ to determine the extent of the overlap (GFP+tdTomato+ dermal cells) between the two models. n = 4 mice.

(B) Representative FACS plots from two replicates as in (A) expressing PDGFRα-H2B-GFP; PDGFRα-CreER; lsl-tdTomato gating first on tdTomato+ cells and then GFP+ to determine the extent of the overlap (GFP+tdTomato+ dermal cells) between the two models. n = 4 mice.

(C) Quantification of the percent overlap of cells expressing both PDGFRα-H2B-GFP and PDGFRα-CreER; LSL-tdTomato, showing an average of 86% overlap regardless of the gating strategy. n= 4 mice p= 0.9098, ns, unpaired, two-sided Welch's t-test.

(D) Quantification of fibroblast depletion in adult mice one week post induction by FACS, comparing PDGFRα-H2B-GFP (WT), PDGFRα-H2B-GFP;lsl-DTA (Control) and PDGFRα-H2B-GFP; PDGFRα-CreER; lsl-DTA (FibDTA) mice. p = 0.4634, ns, *** p = 0.0005, **p = 0.0014. One way ANOVA with multiple comparisons.

(E) Representative FACS plots showing fibroblast heterogeneity in adult mice one week post induction. n=3 control (PDGFRα-H2B-GFP; LSL-DTA) mice compared to n= 3 FibDTA (PDGFRα-H2B-GFP; PDGFRα-CreER ;LSL-DTA) mice after systemic depletion of PDGFRα+ cells. Cells were gated on GFP first, and then gated on CD26+Sca-1-, CD26+Sca-1+, CD26-Sca-1+ and CD26-Sca-1- (Other).

(F) Quantification of fibroblast heterogeneity in adult mice one week post induction. n=3 control (PDGFRα-H2B-GFP; LSL-DTA) mice compared to n=3 FibDTA (PDGFRα-H2B-GFP; PDGFRα-CreER ;LSL-DTA) mice after systemic depletion of PDGFRα+ cells. p = 0.3183 ns (CD26+), p =

0.0524, ns (CD26+Sca-1+), **p=0.01 (CD26-Sca-1+), p=0.3954, ns (CD26-Sca-1-, Other), unpaired, two-sided, Welch's t-test.

(G) Representative whole mount images of control and FibDTA mice adult mice one week post induction after 6hr EdU chase. Epidermal basal cells are represented in magenta and $EdU^+$ cells in white. Scale bar = 50 μm.

(H) Quantification of epidermal basal cell density in control and FibDTA adult mice after systemic tamoxifen induction. Graph represents average number of Hoecsht-positive cells within a 150 $\mu m^2$ area with SD from n = 3 mice. p = 0.2988, unpaired, two-sided, Welch's t-test.

(I) Representative whole mount images of control and FibDTA adult mice 1 week post induction after 6hr EdU chase. Epidermal basal cells are represented in magenta and $EdU^+$ cells in white. Scale bar = 50 μm.

(J) Representative whole mount images of control and FibDTA adult mouse ear tissue 1 week post induction after 6hr EdU chase. Epidermal basal cells are represented in magenta and $EdU^+$ cells in white. Scale bar = 50 μm.

Supplementary Figure 2

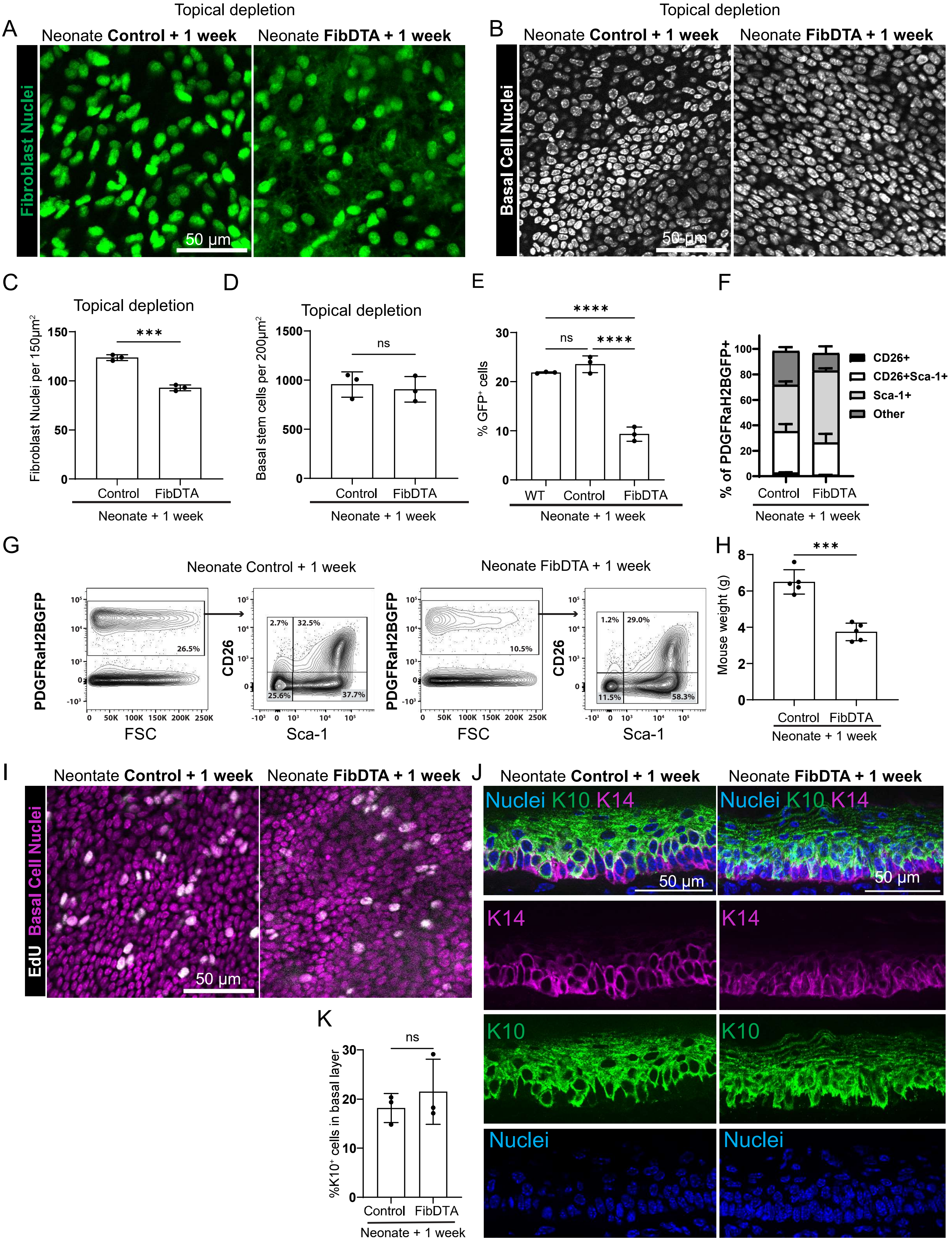

**Supplementary Figure 2.**

(A) Representative images of fibroblast nuclei of control and FibDTA neonatal mice one week following topical administration of 4-OHT. Scale bar = 50 μm.

(B) Representative images of epidermal basal cell nuclei in control and FibDTA mice following topical administration of 4-OHT.

(C) Quantification of fibroblast nuclei identified by the PDGFRα-H2BGFP signal in control and FibDTA neonatal mice one week following topical administration of 4-OHT. n = 3 control and n = 3 FibDTA mice. ***p = 0.0002, unpaired, two-sided, Welch's t-test.

(D) Quantification of epidermal basal cell number in control and FibDTA neonatal mice one week following topical administration of 4-OHT. n= 3 control and 3 FibDTA mice. p = 0.6649, ns, unpaired, two-sided, Welch's t-test.

(E) Quantification of fibroblast depletion in control and FibDTA neonatal mice one week post induction by FACS, comparing PDGFRα-H2B-GFP (WT), PDGFRα-H2B-GFP;lsl-DTA (Control) and PDGFRα-H2B-GFP; PDGFRα-CreER; lsl-DTA (FibDTA) mice. n = 3 mice per group. p = 0.3081, ns, ****p<0.0001. One way ANOVA with multiple comparisons.

(F) Quantification of fibroblast heterogeneity in control and FibDTA neonatal mice one week post induction. n=3 control (PDGFRα-H2B-GFP; LSL-DTA) mice compared to n=3 FibDTA (PDGFRα-H2B-GFP; PDGFRα-CreER ;LSL-DTA) mice after systemic depletion of PDGFRα+ cells. *p = 0.0144 (CD26+), p = 0.2589, ns, (CD26+Sca-1+), **p=0.0014 (CD26-Sca-1+), p=0.0662 ns (CD26-Sca-1-, Other), unpaired, two-sided, Welch's t-test.

(G) Representative FACS plots showing fibroblast heterogeneity in control and FibDTA neonatal mice one week post induction. n=3 control (PDGFRα-H2B-GFP; LSL-DTA) mice compared to n=3 FibDTA (PDGFRα-H2B-GFP; PDGFRα-CreER ;LSL-DTA) mice after systemic depletion of

PDGFRα+ cells. Cells were gated on GFP first, and then gated on CD26+Sca-1-, CD26+Sca-1+, CD26-Sca-1+ and CD26-Sca-1- (Other).

(H) Quantification of mouse weights in n = 5 control and 5 FibDTA neonatal mice one week post induction. *** p = 0.0001 unpaired, two-sided, Welch's t-test.

(I) Representative whole mount images of control and FibDTA neonatal mice one week post induction after 6hr EdU chase. Epidermal basal cells are represented in magenta and $EdU^{+}$ cells in white. Scale bar = 50 μm.

(J) Representative paw tissue sections of control and FibDTA neonatal mice one week post induction stained for keratin10 (K10; green), keratin-14 (K14; magenta) and hoechst to label nuclei (blue). Scale bar = 50 μm.

(K) Quantification for percentage of $K10^{+}$ cells residing in the epidermal basal cell layer in n = 3 Control and n = 3 FibDTA neonatal mice one week post induction.

Supplementary Figure 3

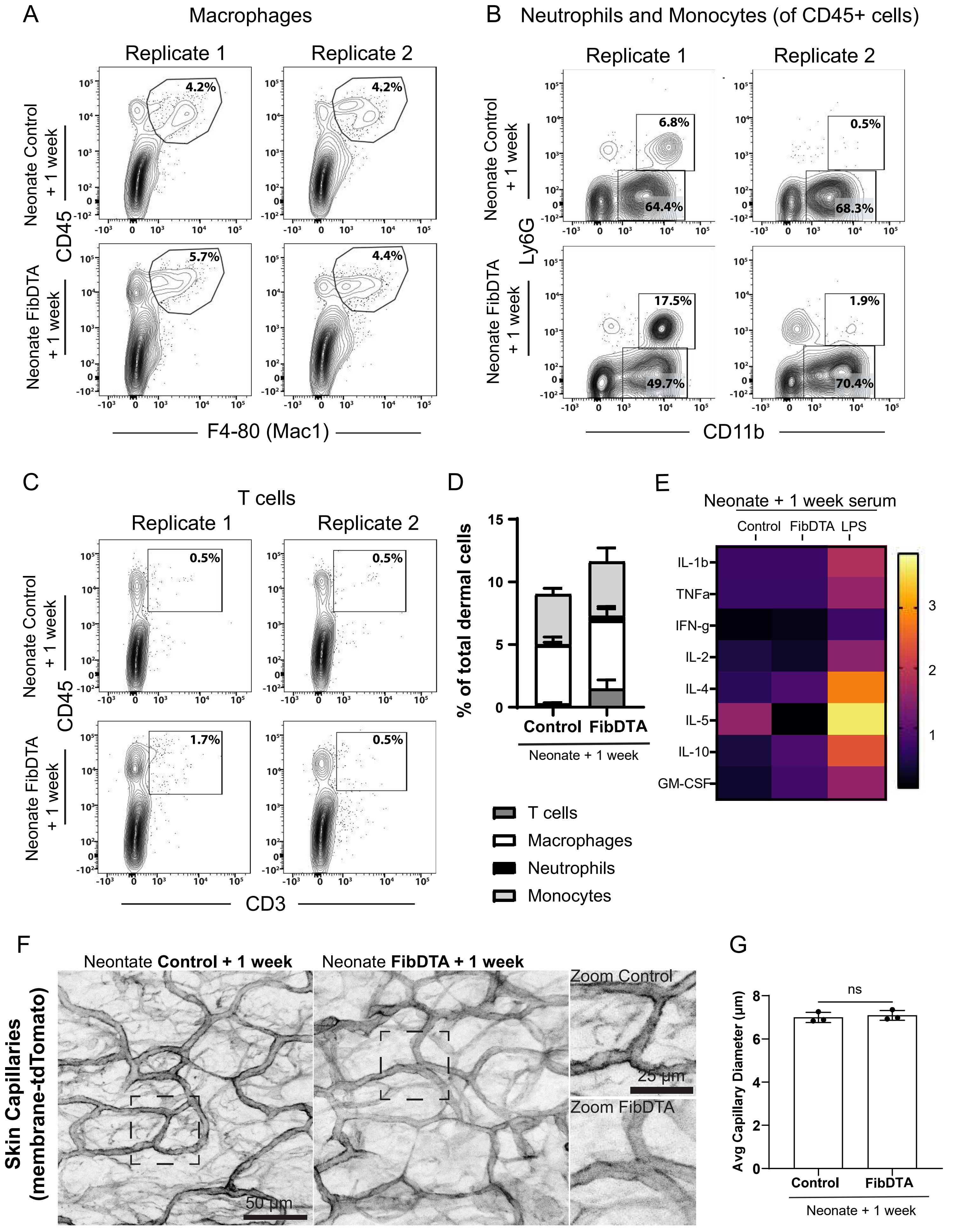

**Supplementary Figure 3.**

(A) Representative flow cytometry plots showing macrophage percentages in dermal preparations of neonatal mice one week post induction from a total of n=3 control (PDGFRα-H2B-GFP; LSL-DTA) mouse paw samples compared to n=4 FibDTA (PDGFRα-H2B-GFP; PDGFRα-CreER ;LSL-DTA) after systemic depletion of PDGFRα+ cells. Cells were gated on CD45 and F4-80.

(B) Representative flow cytometry plots showing neutrophil (Ly6G+Cd11b+) and monocyte (CD11b+) percentages in dermal preparations of neonatal mice one week post induction from a total of n = 3 control (PDGFRα-H2B-GFP; LSL-DTA) mouse paw compared to n=4 FibDTA (PDGFRα-H2B-GFP; PDGFRα-CreER ;LSL-DTA) mice after systemic depletion of PDGFRα+ cells. Cells were gated first on CD45 and then on LY6G and CD11B.

(C) Representative flow cytometry plots showing T cell percentages in dermal preparations of neonatal mice one week post induction from a total of n = 3 control (PDGFRα-H2B-GFP; LSL-DTA) mouse paw compared to n=4 FibDTA (PDGFRα-H2B-GFP; PDGFRα-CreER ;LSL-DTA) after systemic depletion of PDGFRα+ cells. Cells were gated on CD45 and CD3.

(D) Quantification of different immune populations in dermal preparations of neonatal mice one week post induction. n= 3 control (PDGFRα-H2B-GFP; LSL-DTA) mouse paw compared to n=4 FibDTA (PDGFRα-H2B-GFP; PDGFRα-CreER ;LSL-DTA) after systemic depletion of PDGFRα+ cells shown in (A-C). p = 0.2460, ns (macrophages), p = 0.6269, ns (monocytes), p = 0.5126 ns (neutrophils),*p = 0.0448 (T-cells), unpaired, two-sided, Welch's t-test.

(E) Normalized heatmap of cytokine panel taken from serum of neonatal mice one week post induction. n = 3 control, n = 3 FibDTA and n = 2 LPS treated mice. Control vs FibDTA serum p = 0.5545, ns and Control vs LPS serum **p = 0.0066 one way ANOVA with dunnett's multiple comparisons test.

(F) Representative images of the skin capillary plexus, determined by morphology and labeled with LSL-mTmG in control and FibDTA neonatal mice one week post induction. Scale bars = 50 μm and 25 μm for zoom.

(G) Average capillary diameter measurement for n = 3 control and n = 3 FibDTA neonatal mice one week post induction, $p = 0.6436$, ns, unpaired, two-sided, Welch's t test.
